# Molecular detection and characterization of mutation on 23S rRNA gene associated with clarithromycin resistant in *Helicobacter pylori*

**DOI:** 10.1101/650432

**Authors:** Sahar Obi Abd albagi, Hisham Nour aldayem Altayeb, Nadir Musa Khalil Abuzeid

## Abstract

**Background:** *Helicobacter pylori* consider as pathogenic resistant bacterium was colonized mainly in stomach and causing a prolonged gastritis with gastric ulcers were progressing to gastric carcinoma. Also its resistance to antibiotic considered as the main reason for failure to eradicate of this pathogen has been difficult when this resistance occurred as mutant on protein binding site in 23s ribosomal RNA. The highest cure rates have required multidrug antimicrobial therapies including combinations of omeprazole, clarithromycin and metronidazole.

**Result:** Bacterial DNA sequence from gastritis patients with confirmed previous positive ICT samples (Stool and Bloo) were used to obtain co-related between phenotypic & genotypic variant outcome have been observed as SNPs carried on nucleotides which could be altered protein prediction as result of that caused chronic gastritis incline to gastric carcinoma due to abnormal consequence on genetic level in *H. species* (23s rRNA) was referred to clarithromycin resistance, was achieved on this *cross-sectional studies* by running two different primers were amplify in PCR machine, first one for urease producing gene (Glm as universal primer) and second one for 23s rRNA as specific primer (rp1/fp1). Two samples out of Four samples were amplified as final isolate in the first cycle and have a specific band in 23s rRNA (NO.11, NO.24) as further DNA samples investigation were sent to get our target sequence.

**Conclusion:** Bioinformatics tools used to confirm a specific types of mutations give specific position responsible for bacteriostatic activity of macrolides such as clarithromycin depends on capacity to inhibit protein synthesis by binding to the 23S ribosomal subunit (23S rRNA) as resistant gene. The detection tools as MSA (multiple sequence alignment) for our nucleotides sequence to (11&24) samples with Genbank accession number 24_MK208582 and 11_MK208583. One type of mutation has been observed in nucleotide sequence (sample-24) in position 2516 (1344 _complementary) sequence result compared with reference sequence standard reference strain (*H. pylori U27270*) was confirmed which consider it as novel mutation in database for 23S rRNA Gene of *H. pylori* associated with Clarithromycin Resistance gene in Sudanese patients.

## 1. background

Helicobacter pylori (*H. pylori*) one of pathogenic bacteria as stomach organisms of approximately 50% around world’s population, the resistance of *H. pylori* to antibiotics is considered as the main reason to failure eradicate this bacterium ^[1]^. Antibiotic resistance in Helicobacter pylori (*H.pylori*) it’s main factor affecting the efficacy of current treatment methods against the infection caused by this organism, The traditional culture methods for testing bacterial susceptibility to antibiotics are expensive and require 10-14 days, Since resistance to tetracycline, fluoroquinolone, and clarithromycin seems to be exclusively caused by specific mutations in a small region of the responsible gene, molecular methods offer an attractive alternative to the abovementioned techniques; The technique of polymerase chain reaction (PCR) is an accurate and rapid method for the detection of mutations that confer antibiotic resistance ^[2]^.

*H. pylori* is identified as a Group1 carcinogen by the World Health Organization International Agency for Research on Cancer (WHO/IARC), and is associated with the progressed gastric cancer. Eradication of *H. pylori* infection has been reported as an effective strategy in the treatment of peptic ulcers as first line and for any developed gastric mucosa-associated lymphoid tissue lymphoma as well as in the prevention of gastric cancer ^[3]^.

Determining the role of pathogenesis in *Helicobacter pylori*, which involved as gastritis, peptic ulcer disease, gastric cancer, mucosa-associated lymphoid tissue, lymphoma and various extra-gastrointestinal manifestations ^[4–6]^.

In other study the specimen of biopsy like antrum and Corpus biopsies were examined from patients with dyspeptic symptoms *H. pylori* were cultured, and the antimicrobial susceptibility of *H. pylori* was determined using the E-test (clarithromycin, amoxicillin, tetracycline, metronidazole and levofl oxacin) according to the EUCAST breakpoints. Point mutations in the 23S rRNA gene of clarithromycin-resistant strains were investigated using real time PCR ^[7]^.

Some study remained assessed the commercially existing LightMix^®^ RT-PCR analyze for *H. pylori* detection and identification of clarithromycin (CLR) resistance in clinical specimens (gastric biopsies and stool) and culture. The *H. pylori* LightMix^®^ RT-PCR detects a 97 bp long fragment of the 23S rRNA gene and allows the identification of 3 distinct point mutations conferring CLR resistance through melting curve analysis ^[8]^.

Clarithromycin resistance possibly results from the use of the antibiotic in the pediatric, respiratory, and otorhinolaryngology fields ^[9]^. Increase the global clarithromycin-resistance rates from 9% in 1998 to 17.6% in 2008 in Europe also from 7% in 2000 to 27.7% in 2006 in Japan ^[10]^. In patients with clarithromycin-resistant H. pylori, it has been reported that the eradication rate reached with clarithromycin-based regimens shows marked decrease ^[11]^. Infection by gram negative bacillus *(H. pylori)* is more dominant in the developing countries, and more often between younger people reaching up to 10% of the population coverage in comparison to only 0.5% in more developed world The prevalence of *H. pylori* among Sudanese children is 56.3% ^[12]^.

Further study was done in Sudan studied the prevalence of *H. pylori* in Sudanese focuses with gastroduodenal inflammation, *H. pylori* was looked for in biopsy specimens

taken from the antrum by two methods: rapid urease test [Campylobacter-like organism (CLO) test] and culture using Skirrow’s selective supplement.

results of one hundred subjects showed that *H. pylori* was found in 80% of patients with gastritis, 56% of patients with duodenal ulcer, 60% of patients with duodenitis and 16% of normal control subjects ^[13]^. In other common study, the attributed point mutations within the peptidyl transferase-encoding region in domain V of the 23S rRNA gene ^[14,15]^.

Three dissimilar point of mutations have been found associated with macrolide resistance in *H. pylori* strains ^[15,16]^. Mutations A2142G and A2143G are the most frequently reported, whereas mutation A2142C is less common e.g., it was found in 5 out of 129 resistant strains ^[17]^. Two additional mutations, A2115G and G2141A, have been described, occurring in the same strain ^[18]^, but neither has been reported subsequently.

## 2. Objectives

### 2.1 General objective

To detect and characterize mutation in 23S rRNA Gene associated with clarithromycin resistant In H.pylori form Sudanese patients.

### 2.2 Specific Objective

1. To detect causes of chronic gastritis related to gastric carcinoma as result in the patient with resistant H.pylori species by using immunochromatografic Ag/Ab then will be isolate a positive samples to extract bacterial DNA.
2. To co-related between, the progressive symptom in different population coverage associated with Clarithromycin resistant gene.
3. To amplify universal urease producing gene firstly then amplify a target mutant gene (23S rRNA) cause clarithromycin resistant through gene expression technique (PCR).
4. To Analysis the target sequence and determine the type of mutation was observed in NGS as sequencing step for PCR product and applied insilico diagnostic tools on the target sequence of bacterial resistance gene (23S rRNA).

## 3. METHODS

### 3.1 Study design and type of Sampling was obtained for *H.Pylori* strain

This study constructed on descriptive, cross-sectional study.

### 3.2 Setting describe the setting

These simple descriptions for the setting were used

1. Immunochromatografic Antigen and Antibody base test for stool/blood samples.
2. Extracted the DNA by guanidine-HCL method treated sample.
3. *PCR for Amplification of urea producing genes with Glm primers, followed by rp/fp primers as target resistance gene 23S rRNA*.
4. *Gel documentation system*.
5. *Sequencer: sent the positive samples to china “Macrogene companies”*.
6. *bioinformatics tools: Codon Code Aligner “version 8.0.2.6”, Finch TV program “version 1.4.0”*.

#### 3.2.1 Location, relevant date, participant, Sample collection and aerially detection

*Samples were obtained from international Hospital in Khartoum state*, from April *2018-October 2018*. This study includes randomizing Sudanese patients have been infected with *H.pylori* from different age group and both sex of them were included. Set of samples (N= 50) was collected after The clinical report forms (CRFs) were completed by qualified practitioner. Informed consent was obtained from each patient included in the study and the study protocol conforms to the ethical guidelines of the 1975 Declaration of Helsinki as reflected in a priori approval by the Human Ethics Committees of the University of Elnaileen and the Ministry of Health of Sudan. its includes any patients have a symptom of gastritis with *H.pylori* ICT positive Ag/Ab in stool/blood sample consequently. its randomly selected based positive stool sample and blood as sampling technique.

#### 3.2.2 Variables for data source measurement with Diagnostic criteria

##### 3.2.2.1 Preparation of Stool sample and blood sample

Stool and Blood samples were obtained from patients in clean wide and sterile plain containers with immediately diagnosed by ICT Ag for *H.pylori* and then positive samples have been submitted to extract *H.pylori* DNA.

##### 3.2.2.3 DNA extraction from stool and blood samples

Bacterial DNA obtained by using the guanidine-HCL-ethanol supernatant cell after lysed with (SDS) buffer lysis of (WBCs) the cell debris has been removed by centrifuge, then protein was denatured, digested with protease and precipitated with organic solvent such as phenol, and protein precipitated will be removed by centrifuge solution, then discard and isolated pure DNA in sterile (1.5) vacuum tube and preserved in −20°C.

##### 3.2.2.4 Preparation of samples for PCR amplification and further identification Polymerase chain reaction (PCR)

###### Primer design

Firstly, we used two designed primers to each gene (*Glm* and 23s *rRNA*).

The sequence of *Glm-R*(*revers_ AAG-CTT-ACT-TTC-TAA-CAC-TAA-CGC-3*)

*Glm-F*(*forword_AAG-CTT-TTA-GGG-GTG-TTA-GGG-GTT-T-3*) *with PCR product 240bp*.

Primers designated for 23s *rRNA*-f(forward_-*TCG-AAG-GTT-AAG-GAT-GCG-TCA-GTC-3* rRNA-r (reverse:_-*GAC-TCC-ATA-AGA-GCC-AAA-GCC-TTAC-3*_), PCR product given result in 320 bp

###### Preparation of 50 μL PCR master mix

ready PCR tube was containing 10 mM Tris-Cl (pH 9.0), 50 mM KCl, 1.5 mM MgCl, 2.5 U of *Taq* DNA polymerase (Amersham Pharmacia Biotech, Piscataway, N.J.), 200_M deoxynucleoside triphosphate. After diluted each revers and forward primers of (*Glm* and *23sr RNA*) sterile water at 10:9 microliters in separated PCR tubes, then Samples were taken 1μl + 17μl from D. W+ 1 μl of extracted bacterial DNA were added to iNtRON Master mix tubes. See details of Table 1.

###### PCR program

Amplification will be carried out in a DNA thermal cycler (model 9700; Perkin-Elmer Corporation, Norwalk, Conn.).

###### Programed cycle of *Glm* primer

Thirty cycle, each 30s at 95°C, 30s at 64.1°C, 40.0s at 72°C, were performed 2 min of denaturation at 95°C. cycle followed by final elongation at 72°C for 5min.

###### Programed cycle of 23s rRNA primer

Thirty-five cycles, each 15 s at 95°C, 20 min at 60°C, and 30s at 68°C, were performed after 5 min of denaturation at 95°C. Cycles will be followed by a final elongation at 68°C for 2 min.

##### 3.2.2.5 Gel Electrophoresis

Fore microliter portions of the PCR products will be analyzed by electrophoresis in 2% gel agarose within a 1.5% agarose gel in Tris-acetate-EDTA (TAE) buffer stained with ethidium bromide in parallel with a molecular weight marker: Gene ruler 100-bp DNA ladder (MBI Fermentas; Vilnius, Lithuania), (as trial steps) then Steps procedure were compiled for another remaining samples. Then Specific bands were observed result as under gel documentation machine after ladder was running and separated have specific pieces for every 100 bp.

##### 3.2.2.6 Sequencing process and bioinformatics tools

The PCR product give a positive result at both genes produced specific bands, as complementary steps we send products to Global Genomic Services company (**BGI**) in china to sequencing positive product and provide good quality genomic data.

In the sequencing primers **Rp-1** 23s *rRNA*-f(forward_*TCG-AAG-GTT-AAG-GAT-GCG-TCA-GTC-3*rRNA-r **Rp-2** (reverse:_ *GAC-TCC-ATA-AGA-GCC-AAA-GCC-TTAC-3*_), with PCR product will give result in 320 bp. (GenBank accession number of ref sequence U27270). After sequencing process, bioinformatics tools were used to detect specific type of mutant gene which was located in specific position was responsible for bacteriostatic activity of macrolides such as erythromycin and clarithromycin depends on the capacity to inhibit protein synthesis by binding to the 23S ribosomal subunit (23S rRNA) as resistant gene. For that reason, *in silico* diagnostic tests were applied as new promising approach to study type of mutant for desired gene (23S rRNA) caused drugsusceptibility.

Our nucleotide sequencing result data with extra comparison sequences were download from NCBI was analyzed by using alignment program of Codon Code Aligner “version 8.0.2.6” includes phylogeny and BLAST online programs. Also we observed the signal of nucleotide mutant by using Finch TV program “version 1.4.0”, *In silico* tools were used these as collaborator programs as enhancement analysis of our target result.

**Bias:** N.A

**Study size:**

*this study includes fifty (N= 50) samples from diagnosed population then isolate a Helicobacter* pylori positive in stool and blood sample, (N=Z*2 *P*Q/D2).

**Statistical methods:** N.A

## 4. Result

### 4.1 Immunochromatographic ICT Ag/Ab detection analysis result

Out of 50 different samples (25blood/25stool) were yielded positive ICT samples total “15 from stool/ 6 samples from blood” positive with *H.pylori* positive respectively then collect it for further investigation.

### 4.2 PCR amplification of *Glm* and sequence analysis

Primers of *Glm-F* and *Glm-R*, designed to amplify Urea producing gene fragment From *H. pylori*, yielded a 240-bp PCR product. Amplicons were achieved to all the positive ICT-Ag/Ab base were obtain 21 *H. pylori* positive extracted samples examined. To scan the genotypic method according to our positive samples at both kinds of it, we consider *Glm* as a universal primer to provide a subsequent step to detect mutation in 23sr rRNA as gene resistance of clarithromycin.

Our result shows positive for 240bp in four samples “11,24,13,15,16” there are having our universal desired gene of *Glm*. Only 4 out of 5 were selected in next amplification due to poorly observed band of sample 16.

Fig.1 illustrates the digestion gel electrophori’s resulting in documentation system of 5 positive samples.

**Figure 1:**
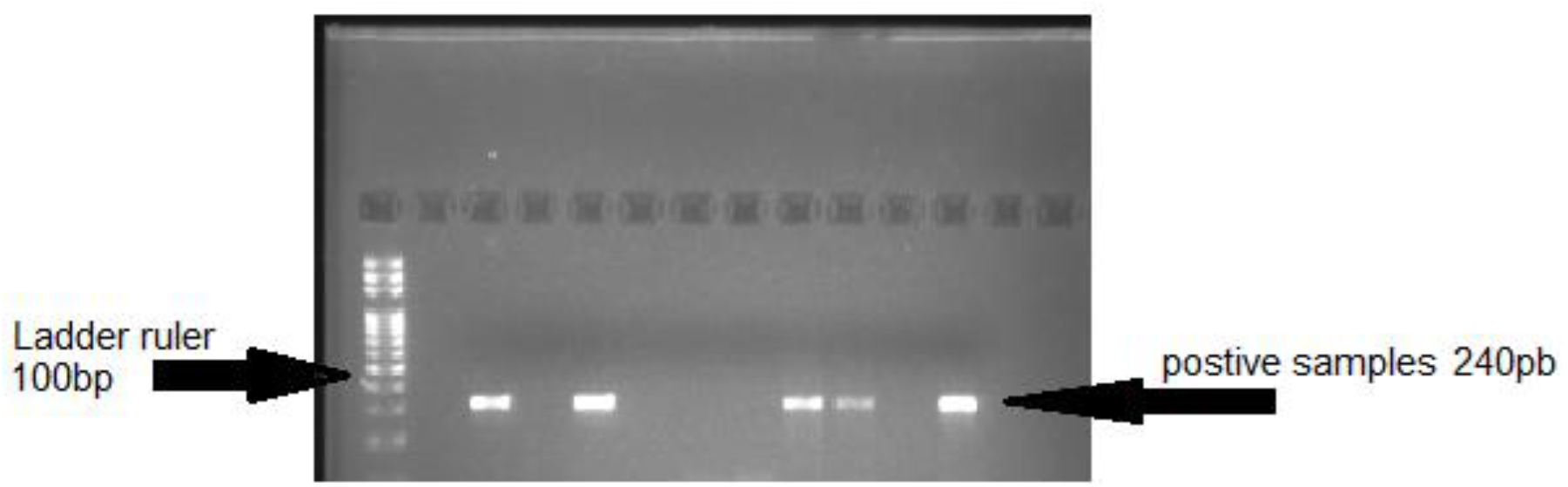
Glm positive samples “11,13,16,15,24” with 240-pb of urease producing gene. Ladder fragment length is 100bp. When no bands are observed in remaining samples.

### 4.3 PCR amplification of 23S rRNA and SNPs sequence analysis

Primers Fp-1 and Rp-1, designed to amplify 23S rRNA fragment from *H. pylori*, yielded a 320-bp PCR product. Amplicons were obtained for all the 4 *H. pylori Glm* positive samples were examined. To examine the genetic basis as a genotypic method to confer a positive universal primer (*Glm*; urease producing gene) of previous ICT positive in phenotypic method, we analyzed the nucleotide sequence, for all the resistant isolates, of the 320-bp PCR product of the 23S rRNA gene, and we compared these sequences with those obtained for the standard reference strain (*H. pylori U27270*) as well as with selected positive isolate DNA sample in *Glm*. No one of isolates showed the mutation defined at positions 2142 and 2143 ^[19, 20]^. Particularly, Fig.2 shown illustrates the digestion gel electrophori’s resulting in documentation system. the sequence analysis of two resistant isolates when its showing a specific band in both gene resistance of clarithromycin (23s rRNA), presented a single point mutation: the transversion from T to C at position 1344 in complementary and from G to A in position of 2516. the nucleotide sequence of reverse and forward resistant strain was being reported, while in Fig.3 was showed mutant signal by Finch TV. Ink program version 1,4.0 of the same samples an arrow the position of T1344C mutation in the model of the 23S rRNA. The remaining isolates, for which positive 23sr RNA, did not existence of a T1344C mutation. Finally, we use Cod Codon Align program version 8.0.2.6, to observe SNP and comparing it with reference sequence line group in phylogeny tree in Fig 5,6.

**Figure 2:**
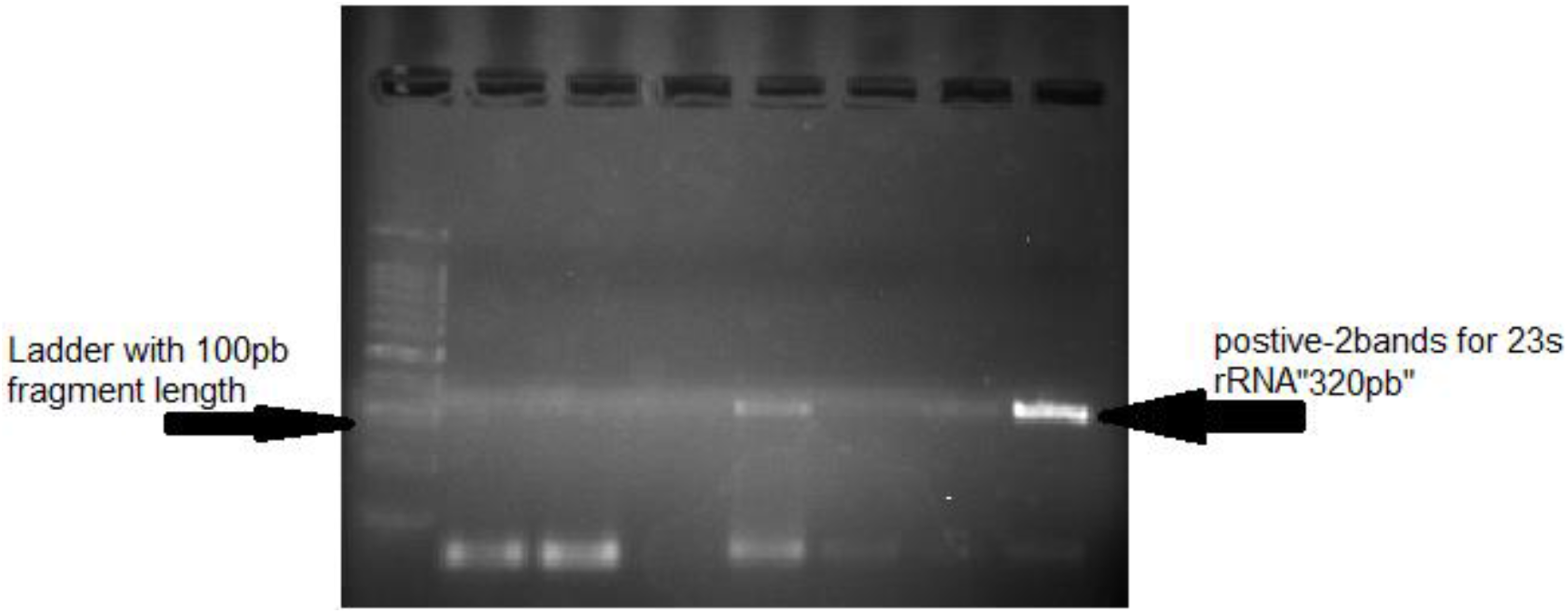
digestion gel electrophori’s result of 23srRNA for “11,24” were observe specific bands in “320pb”. When no bands are observed in “13, 15, 16” sample, with size marker 100bp.

**Figure 3:**
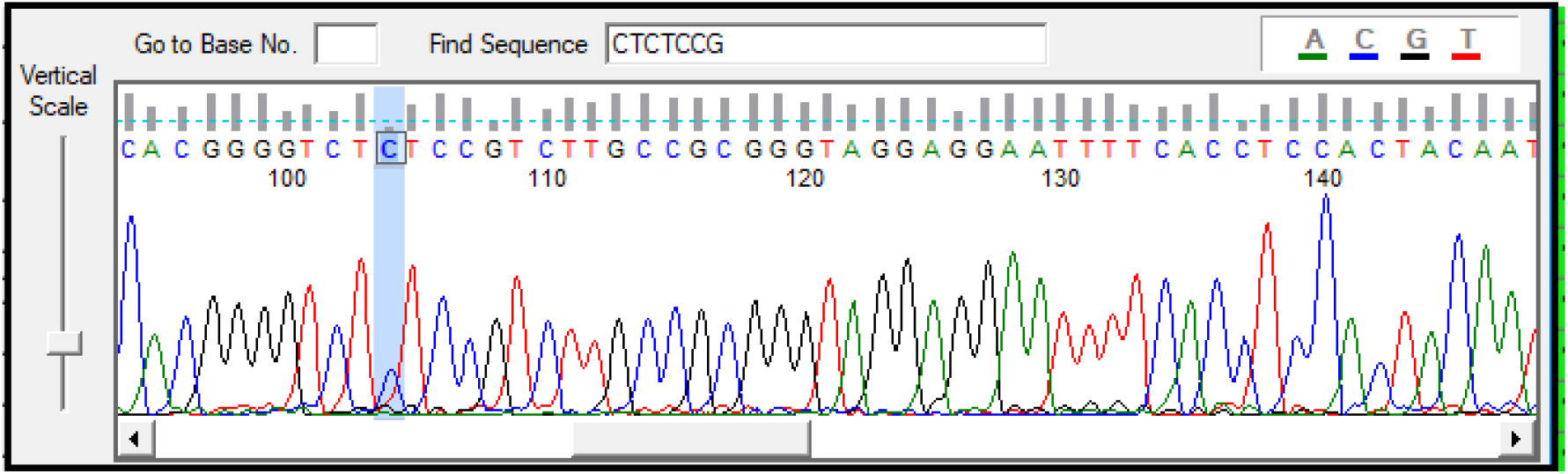
sample_11 forward sequence show point mutation signal T ➔ C (Thr ➔ Cys). When signal was observed with low graphic.

**Figure 4:**
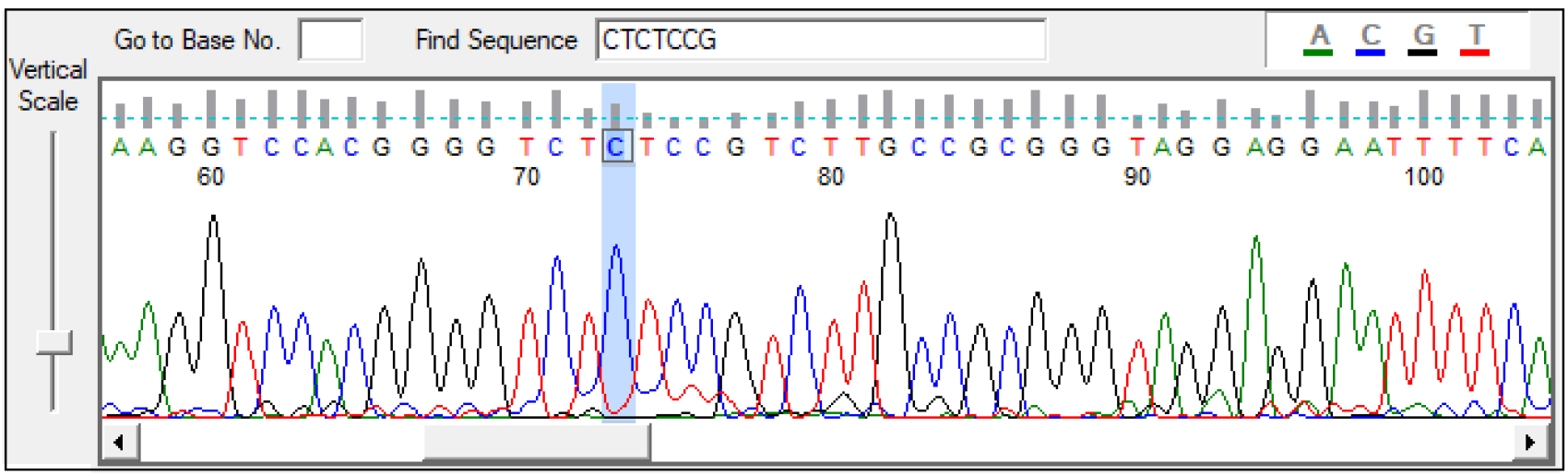
sample_11 revers sequence show point mutation signal T ➔ C (Thr ➔ Cys). When signal was observed with clear large sign. illustrate by Finch TV (1.4.0 version).

**Figure 5:**
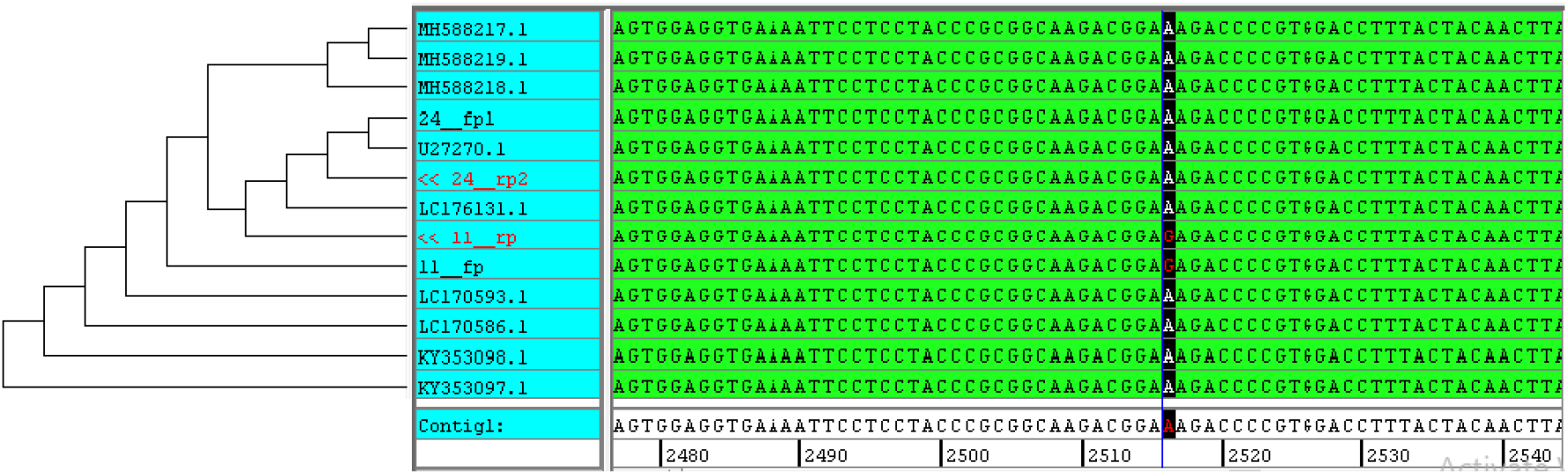
Phylogeny tree with MSA of forward coting sequences compared with ref sequence “U27270.1”, was show SNPs on position 2516 in Query samples “no.11_reveres/forward”.

**Figure 6:**
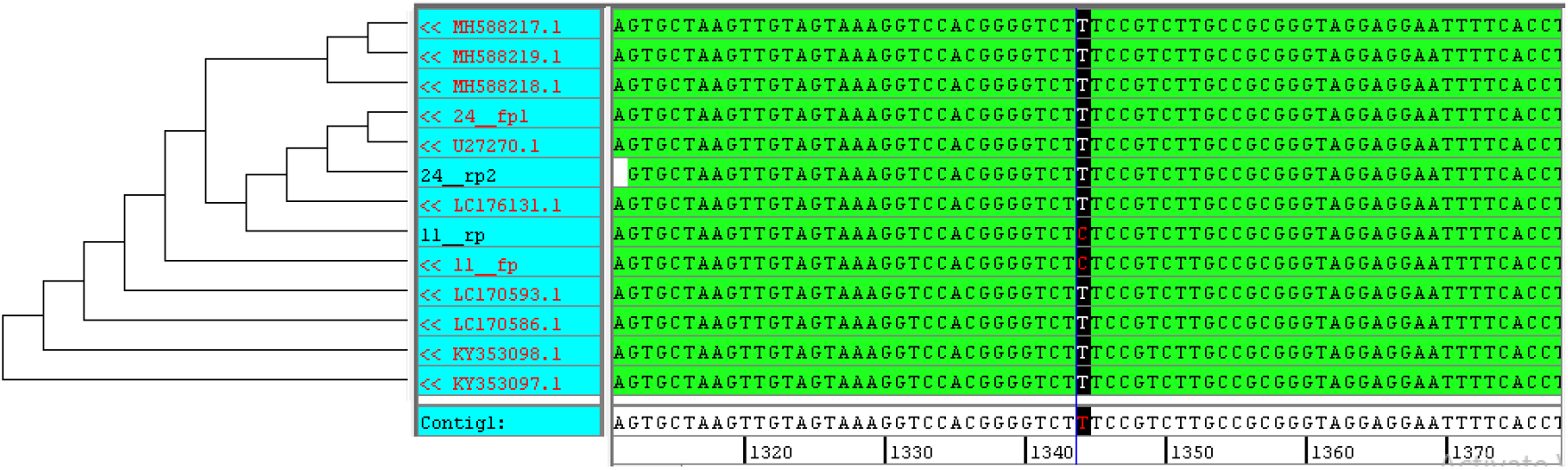
phylogeny tree with MSA of revers, was show SNPs on position 1344 in Query samples “no.11_reveres/forward”.

## 5. Discussion

### Key results, limitation and interpretation

In this study, meanly focusing on genotypic methods to obtain co*-related between* phenotypic-genotypic result variation have been observed the mutant type carrier on nucleotide sequences *as result of an* effect on the genetic level of *H.pylori species with* (23s rRNA) in clarithromycin resistance *in different population associate*.

- Accordingly, positive *immunochromatografic* antigen results of 21 positives from 50 collected samples were includes equal samples (stool\blood), depending on previous questioner. Extraction of H*.pylori* DNA samples were excluded from 21 positive “Ag-Ab ICT samples” using the guanidine-HCL-ethanol method.
- The amplification of urease producing genes were using universal genes Glm primers (forward and reverse) after run in Gel-electrophoresis and observed specific bands “240-pb” in Gel documentation system, so according to product length of target gene and positive control with product length of 240 bp. Five isolates samples have the same specific bands were described in manual of desired primer was show in trial of control positive, sample “no.16” were exclude due to weekly appearance of band. The negative result of remaining samples in PCR were referred to low specify in quality of ICT devices mainly in Ab detection, it was predicting nonspecific antibody which resulting in more cross matching with others similar antibody’s producing diseases.
- Preserved extracted samples giving positive in urea gene were amplified in second one targeting gene responsible for clarithromycin resistance gene 23s rRNA, for both primers (Fp-1 and Rp-1) Amplifications were carried out in a DNA thermal cycle resulting with observe our result under UV light of gel documentation with two specific bands at 320bp but others three remaining sample have been no result at 320bp. That negative result in universal primer may be due to two reasons either have no desired DNA in our samples or have some bacterial strain doesn’t longer to *H.pylori* strains. The result was abbreviating in Table 2.
- Finally, some detection tools in our received sequence of “no.11 and no.24”. One type of mutation in sample-11 reverse and forward nucleotide sequence in our observed result have a different mutant nucleotide locution from database sequences was observed new position in 2516 and in 1344 of complementary results. And we consider it as novel mutation in the 23S rRNA Gene of Helicobacter pylori was associated with Clarithromycin Resistance gene in Sudanese patients these nucleotide sequence was available in GenBank after direct submission of sequence data with GenBank accession numbers is SUB4631272 24 MK208582, SUB4631272 11 MK208583.
- Follow these link respectively https://www.ncbi.nlm.nih.gov/nuccore/MK208582 https://www.ncbi.nlm.nih.gov/nuccore/MK208583.1
- The signal of mutant nucleotide in sample 11 has good reflection in signal sign in Fig.3,4 of Finch program was indicted to real substitution from T to C. Also multiple alignment programming tool of Codon code align were used for both sequence “11,24” to detect whose one closes to reference sequence standard reference strain is (*H. pylori U27270*) and was seen sample-24 it’s more closes to the reference sequence U27270.1 group and LC176131.1, this sequence report additional mutation at T1958G in resistant strain of clarithromycin ^[21]^, but sample-11 appears in different line group as the phylogeny tree result support were referred that to single mutant nucleotide and its more nearly to MH588218.1 strain “OC975” of 23S ribosomal RNA gene sequence as seen both of them separated from the same point so it’s more likely to be more closes reading in multiple sequence alignment then subsequently possibly to have mutant type gives the same protein prediction of this strain sequence. For Phylogeny illustrate See Fig.5,6.

## 6. Conclusions

23s rRNA gene related to some SNPs responsible to resist in clarithromycin were detected by using different methods; immunochromatographic test, PCR for two possible related genes and sequencer to get linear nucleotide sequence then analyzed it with using bioinformatics prediction tools to check the type of the mutation on the nucleotides sequence and determination of the differences between database sequence with our target sequence using phylogeny tool. One SNPs were detected to be possible affected on Helicobacter pylori resistance.

## 7. Declarations

### Authors’ contributions

All authors support to gets final designed primer, also practical helper and reviewer the manuscript. The authors read and approved the final manuscript.

## Acknowledgements

First of all, we would like to express my special thanks to Allah who gave me the strength and patient to accomplish this work. I would like also to express my appreciation and thanks to Dr. Nadir Musa Khalil Abuzeid department of Medical Microbiology, Assistant Professor and Consultant of Medical Microbiology Also Dr. Hisham Nouraldayem Altayeb Mohamed Assistant professor of molecular biology, and teacher in research molecular lab of AL-neelain University, whose supported me with valuable help and encouragement throughout the period of this study. I never forget first person who was and still is a base of support and motivate thanks Mom.

## Availability of data and materials

The datasets used and/or analyzed during the current study are available from the corresponding author on reasonable request.

## Consent for publication

Not applicable.

## Ethics approval and consent to participate

The clinical report forms (CRFs) were completed by qualified practitioner. Informed consent was obtained from each patient included in the study and the study protocol conforms to the ethical guidelines of the 1975 Declaration of Helsinki as reflected in a priori approval by the Human Ethics Committees of the University of Al Naileen and the Ministry of Health of Sudan. This to certify this reasearch project under title: Molecular detection and characterization of mutation on 23S rRNA gene associated with clarithromycin resistant in *Helicobacter pylori* 2018.

Prapered and submitted by: Sahar Obi Abd Albagi Mohammed.

Hase been approved by the institutional review board, Al Neelain University **IBR Serial No: NU-IBR-19-2-2-15.**

## 8 Abbreviations

rRNA: ribosomal RNA
MSA: Multiple Sequence Alignment
NGS: Next Generation Sequencing
BLAST: Basic Local Alignment Searching Tools
NCBI: National Center for Biotechnology Information.
CLR: Clarithromycin
*Glm*: primer referred to urease producing gene.
320-bp: 320-base pair.
PCR: Polymerase Chain Reaction.
Fp-1 and Rp-1: Forward and Reverse primers for Clarithromycin Resistance Gene. (N=Z*2 *P*Q/D2): This Equation of sample size, Whereby: N= sample size, Z= stander deviation, P= frequency of occurrences, Q= frequency of non-occurrences, D= level of precision.

## REFERENCES

[1] Khademi F, Poursina F, Hosseini E, Akbari M, Safaei HG. Helicobacter pylori in Iran: A systematic review on the antibiotic resistance. Iran J Basic Med Sci. 2015 Jan;18(1):2–7. PubMed PMID: 25810869; PubMed Central PMCID: PMC4366738.

[2] Nishizawa T, Suzuki H. Mechanisms of Helicobacter pylori antibiotic resistance and molecular testing. Front Mol Biosci. 2014;1:19. Published 2014 Oct 24. doi:10.3389/fmolb.2014.00019.

[3] Fukase K., Kato M., Kikuchi S., Inoue K., Uemura N., Okamoto S., et al (2008) Effect of eradication of Helicobacter pylori on incidence of metachronous gastritis carcinoma after endoscope resection of early gastric. Lancet. 2008 Aug 2; 372(9636): 392–397. doi: 10.1016/S0140-6736(08)61159-9

[4] Georgopoulos SD, Papastergiou V, Karatapanis S. Current options for the treatment of Helicobacter pylori. Expert Opin Pharmacother 2013; 14:211–223.

[5] Aydin A, Oruc N, Turan I, Ozutemiz O, Tuncyurek M, Musoglu A. The modified sequential treatment regimen containing levofl oxacin for Helicobacter pylori eradication in Turkey. Helicobacter 2012; 14:520–524.

[6]. Cuadrado-Lavín A, Salcines-Caviedes JR, Carrascosa MF, Dierssen-Sotos T, Cobo M, Campos MR, et al. Levofl oxacinversus clarithromycin iin a 10day triple therapy regmen for fiirstline Helicobacter pylori eradication: a single-blind randomized clinical trial. J Antimicrob Chemother 2012; 67: 2254–2259

[7]. Caliskan, R., Tokman, H.B., Erzin, Y., Saribas, S., Yuksel, P., Bolek, B.K., Sevuk, E.O., Demirci, M., Yilmazli, O., Akgul, O. and Kalayci, F., 2015. Antimicrobial resistance of Helicobacter pylori strains to five antibiotics, including levofloxacin, in Northwestern Turkey. Revista da Sociedade Brasileira de Medicina Tropical, 48(3), pp.278–284.

[8]. Redondo J. Jareño., M. Keller P., Zbinden R., Karoline Wagner J.J. Redondo et al. / Diagnostic Microbiology and Infectious Disease 90 (2018) 1–6

[9]. Kaneko, F., Suzuki, H., Hasegawa, N., Kurabayshi, K., Saito, H., Otani, S., Nakamizo, H., Kawata, K., Miyairi, M., Ishii, K. and Ishii, H., 2004. High prevalence rate of Helicobacter pylori resistance to clarithromycin during long-term multiple antibiotic therapy for chronic respiratory disease caused by non-tuberculous mycobacteria. Alimentary pharmacology &therapeutics, 20, pp.62–67.

[10]. Asaka M., Kato M., Takahashi S., Fukuda Y., Sugiyama T., Ota H., etal. (2010). Guide lines for the managemen to f Helicobacterpylori infection inJapan:2009 revisededition. Helicobacter 15, 1–20. doi:10.1111/j.1523-5378.2009.00738.x

[11]. Nishizawa T., Maekawa T., Watanabe, N., Harada, N., Hosoda, Y., Yoshinaga, M., et al. (2014). Clarithromycin Versus Metronidazole as First-line Helicobacter pylori Eradication ?amulti center, prospective, randomized controlled study in Japan. J. Clin. Gastroenterol, doi: 10.1097/MCG.0000000000000165. [Epub ahead of print].

[12]. Salih, K.M., Elfaki, O.A., Hamid, Y.H., Eldouch, W.M., Diab, M. and Abdelgadir, S.O., 2017. Prevalence of Helicobacter Pylori among Sudanese children admitted to a specialized children hospital. Sudanese journal of paediatrics, 17(1), p.14.

[13]. Azim Mirghani YA, Ahmed S, Ahmed M, Ismail MO, Fedail SS, Kamel M, Saidia H (1994). Detection of *Helicobacter pylori* in endoscopic biopsies in Sudan: Trop Doct; 24(4):161–3.

[14]. Taylor, D. E., Z. Ge, D. Purych, T. Lo, and K. Hiratsuka. 1997. Cloning and sequence analysis of two copies of a 23S rRNA gene from *Helicobacter pylori* and association of clarithromycin resistance with 23S rRNA mutations. Antimicrob. Agents Chemother. 41:2621–2628.

[15]. Versalovic, J., D. Shortridge, K. Kibler, M. V. Griffy, J. Beyer, R. K. Flamm, S.K. Tanaka, D. Y. Graham, and M. F. Go. 1996. Mutations in 23S rRNA are associated with clarithromycin resistance in *Helicobacter pylori*. Antimicrob. Agents Chemother. 40:477–480.

[16]. Versalovic, J., M. S. Osato, K. Spakovsky, M. P. Dore, and R. Reddy. 1997. Point mutations in the 23S rRNA gene of *Helicobacter pylori* associated with different levels of clarithromycin resistance. J. Antimicrob. Chemother. 40: 283–286.

[17]. van Doorn, L. J., Y. Glupczynski, J. G. Kusters, F. Megraud, P. Midolo, N. Maggi Solca, D. M. M. Queiroz, N. Nouhan, E. Stet, and W. G. V. Quint. 2001. Accurate prediction of macrolide resistance in *Helicobacter pylori* by a PCR line probe assay for detection of mutations in the 23S rRNA gene: multicenter validation study. Antimicrob. Agents Chemother. 45:1500–1504.

[18]. Hultén, K., A. Gibreel, O. Skold, and L. Engstrand. 1997. Macrolide resistance in *Helicobacter pylori:* mechanism and stability in strains from clarithromycin-treated patients. Antimicrob. Agents Chemother. 41:2550–2553

[19]. Debets-Ossenkopp Y. J., M. Sparrius, J. G. Kusters, J. J. Kolkmann, and C. M. J. E. Vandenbroucke-Grauls. 1996. Mechanism of clarithromycin resistance in clinical isolates of *Helicobacter pylori*. FEMS Microbiol. Lett. 142:37–42.

[20]. Versalovic, J., M. S. Osato, K. Spakovsky, M. P. Dore, R. Reddy, G. G. Stone, D. Shortridge, R. K. Flamm, S. K. Tanaka, and D. Y. Graham. 1997. Point mutations in the 23S rRNA gene of *Helicobacter pylori* associated with different levels of clarithromycin resistance. Antimicrob. Agents Chemother. 40:283–286.

[21]. Miftahussurur, M., Cruz, M., Subsomwong, P., Abreu, J.A.J., Hosking, C., Nagashima, H., Akada, J. and Yamaoka, Y., 2017. Clarithromycin-based triple therapy is still useful as an initial treatment for Helicobacter pylori infection in the Dominican Republic. The American journal of tropical medicine and hygiene, 96(5), pp.1050–1059.

